# Vestibular Compound Action Potentials and Macular Velocity Evoked by Sound and Vibration in the Guinea Pig

**DOI:** 10.1101/2022.09.29.510190

**Authors:** Christopher J. Pastras, Ian S. Curthoys, Richard D. Rabbitt, Daniel J. Brown

## Abstract

To examine mechanisms responsible for vestibular afferent sensitivity to transient air conducted sounds (ACS) and inter-aural bone conducted vibration (BCV), we performed simultaneous measurements of stimulus-evoked vestibular compound action potentials (vCAPs), utricular macula or stapes velocity, and Vestibular Microphonics (VMs) in the anaesthetized guinea pig. For short duration punctate stimuli (<1ms), the vCAP increases magnitude in close proportion to macular velocity and temporal bone (ear-bar) acceleration, rather than other kinematic variables. For longer duration stimuli, the vCAP magnitude switches from acceleration sensitive to linear jerk sensitive. vCAP input-output (IO) functions suggest primary afferent response generation has the same origins for both BCV and ACS, with similar macular velocity thresholds and IO functions for both stimuli. Frequency tuning curves evoked by tone-burst stimuli also show the vCAP increases magnitude in proportion to macular velocity, while in contrast, the VM increases magnitude in proportion to macular displacement across the entire frequency bandwidth tested. The subset of vestibular afferent neurons responsible for synchronized firing and vCAPs make calyceal synaptic contacts with type I hair cells in the striolar region of the epithelium and have irregularly spaced inter-spike intervals at rest. Present results provide new insight into mechanical and neural mechanisms underlying synchronized action potentials in these sensitive afferents, with clinical relevance for understanding the activation and tuning of neurons responsible for driving rapid compensatory reflex responses.

**Significant statement:** Calyx-bearing afferents in the utricle have the remarkable ability to fire an action potential at a precise time following the onset of a transient stimulus and provide temporal information required for compensatory vestibular reflex circuits, but specifically how transient high-frequency stimuli lead to mechanical activation of hair cells and neural responses is poorly understood. Here, we dissect the relative contributions of mechanics, hair cell transduction, and action potential generation on short-latency responses to transient stimuli. Results provide a framework for the interpretation of synchronized vestibular afferent responses, with relevance to understanding origins of myogenic reflex responses commonly used in the clinic to assay vestibular function, and vestibular short latency potentials commonly used for vestibular phenotyping in rodents.

## Introduction

Vestibular otolith organs are phylogenetically ancient inertial sensors that evolved hundreds of millions of years ago in primitive fish ^1^, and successfully endowed extant land-dwelling vertebrates with the sensory neural inputs necessary for locomotion and navigation in a complex terrestrial environment ^2–5^. In amniotes, some otolith afferent neurons preferentially respond to low-frequency gravito-inertial acceleration ^6–11^, while others preferentially respond to high-frequency air conducted sound (ACS) or bone conducted vibration (BCV) ^12–19^. The full population of otolith sensory neurons provide the central nervous system with broad-band detection of linear acceleration and head orientation in three-dimensional (3D) space, providing critical inputs to the autonomic nervous system to modulate heart rate and respiration during movements ^20,21^, and to motor circuits responsible for the vestibular-ocular, -spinal, and -colic reflexes ^9,22^. The compensatory nature of vestibular circuits makes disorders of the otolith organs particularly debilitating, often leading to sensory conflict and symptoms of dizziness, nausea, blurred vision, anxiety, and disorientation. Otolith function is commonly tested in the clinic using transient ACS or BCV to evoke reflexive cervical or ocular myogenic potentials (VEMPs), but precisely how high-frequency transient stimuli lead to mechano-transduction and neural responses in otolith organs is not well understood.

The broad dynamic range of otolith sensitivity from DC to several kilohertz 23 arises from diverse properties of hair cells, synapses, and vestibular afferent spike generators ^24,25^. Amniote neuroepithelia have two major hair cell types (I and II) and two major synaptic terminal types (bouton, calyx, or their combination; dimorphic)^26–28^ which combine to provide the broad frequency bandwidth and diversity in action potential generation between different afferent neurons. The larger diameter calyx bearing afferents, which evolved in land-dwelling amniotes ^2,3,29^, make synaptic contacts with type-I hair cells in the striolar region of the macula ^30–32^, and are characterized by their irregular action potential discharge rate, phasic responses to maintained stimuli, and sensitive short-latency responses to linear acceleration^33^. Calyx synaptic terminals completely envelop one or more type-I hair cells and are exquisitely sensitive to transient stimuli ^18,34^. Three modes of excitatory synaptic transmission occur at calyx terminals: quantal glutamatergic vesicular release (QT) ^35,36^, ultrafast nonquantal ephaptic coupling (NQf) ^24,37,38^, and slow nonquantal accumulation of K^+^ within the synaptic cleft (NQs) ^24,39,40^. Direct ephaptic electrical coupling (NQf) is the component responsible for ultrashort latency and high sensitivity of calyx bearing vestibular afferents to transient inputs ^37,38^.

Sensitivity of calyx bearing otolith afferent neurons to transient ACS and BCV is routinely exploited in the clinic and the laboratory to test otolith function. In the clinic, reflexive cervical and ocular vestibular evoked myogenic potentials (cVEMP and oVEMP) are used to test saccular and utricular function ^41^, and in the laboratory short latency vestibular stimulus evoked potentials (VsEP) are used to screen otolith function in mice and other rodents ^42^. VsEPs are compound action potentials arising from transient stimuli that evoke nearly synchronous firing of a large number of calyx-bearing afferent neurons. When the vestibular compound action potential (vCAP) is recorded from localized sites near the vestibular nerve branch such as the facial nerve canal, the signal-to-noise ratio is enhanced providing recordings similar to auditory CAPs recorded from the round window niche ^43,44^. vCAPs reflect combined responses of the population of sensitive afferent neurons and have been recorded in both acute and chronic animal models of health and disease^45^. Although whole-nerve neural responses to transient ACS and BCV stimuli have been reported for otolith organs, it is currently not known how high frequency transient stimuli lead to mechano-electrical transduction (MET) by sensory hair cells or the generation of synchronized action potentials.

The present report quantifies the relationship between ACS and BCV stimuli, mechanical vibration of the macula, gating of hair cell MET channels, and generation of vCAPs in the guinea pig utricle. This was achieved by simultaneous measurement of temporal bone acceleration, macular (and stapes) velocity, vestibular microphonics (VMs), and extracellular vCAPs. Results quantify the relationship between mammalian utricular mechanics, VMs and vCAPs evoked by ACS and BCV, as well as the timing and sensitivity of primary afferent signals driving compensatory vestibular reflexes to transient stimuli.

## Methods

### Animal preparation & surgery

Experiments were performed on 28 adult tri-colored guinea pigs (*Cavia porcellus*) weighing between 300-500g of either sex. This study was carried out in accordance with the recommendations of the University of Sydney Animal Care and Ethics Committee (Approved protocol number: 2019/1533). Prior to procedures, animals first received pre-anesthetic intraperitoneal injections of Atropine Sulphate; 0.1mg/kg (0.6mg/ml; Apex Laboratories, NSW, Australia) and Buprenorphine Hydrochloride; 0.05mg/kg (Temgesic; 324μg/ml; Reckitt Benckiser, Auckland, NZ). Thereafter, animals were anesthetized in an induction chamber with Isoflurane (2-4%; Henry Schein, NSW, Australia) saturated in medical O_2_ (Coregas, NSW, Australia). Once lacking a foot-withdrawal reflex, guinea pigs were transferred to the surgical table, and received anesthetic via a nose cone, whilst local injections of lignocaine hydrochloride (Lidocaine, Troy Laboratories, NSW, Australia) were delivered to surgery sites. Animals were then tracheotomized and artificially ventilated using Isoflurane (~2%) with oxygen, with the aid of a small animal ventilator (Model 683, Harvard Apparatus, MA, USA). Animals were thereafter rigidly mounted in custom-made ear-bar frames, housing a ‘canalphone’ speaker (ATH-IM70, Audio-Technica, Tokyo, Japan) predominately used for broadband masking and high-frequency stimuli. (Note, high-frequency with regards to the vestibular system, being between 1-2kHz, which is considered low-frequency for the mammalian cochlea). Low-frequency air-conducted sound was delivered to the ear-canal via a silastic tube sealed into the earbar opening, which was coupled to a modified ‘sub-woofer’ speaker (Beyerdynamic, Heilbronn, Germany) acting as a volume velocity source. For sound pressure level calibration, a low-noise microphone probe (ER10B, Etymotic Inc., IL, USA) was sealed into the earbar opening. For the delivery of vibrational stimuli, an electrodynamic mini-shaker (Type-4810, Brüel & Kjær, Denmark) was attached to the earbar in the inter-aural plane via a 5cm metal rod (Fig. 1).

**Fig. 1.**
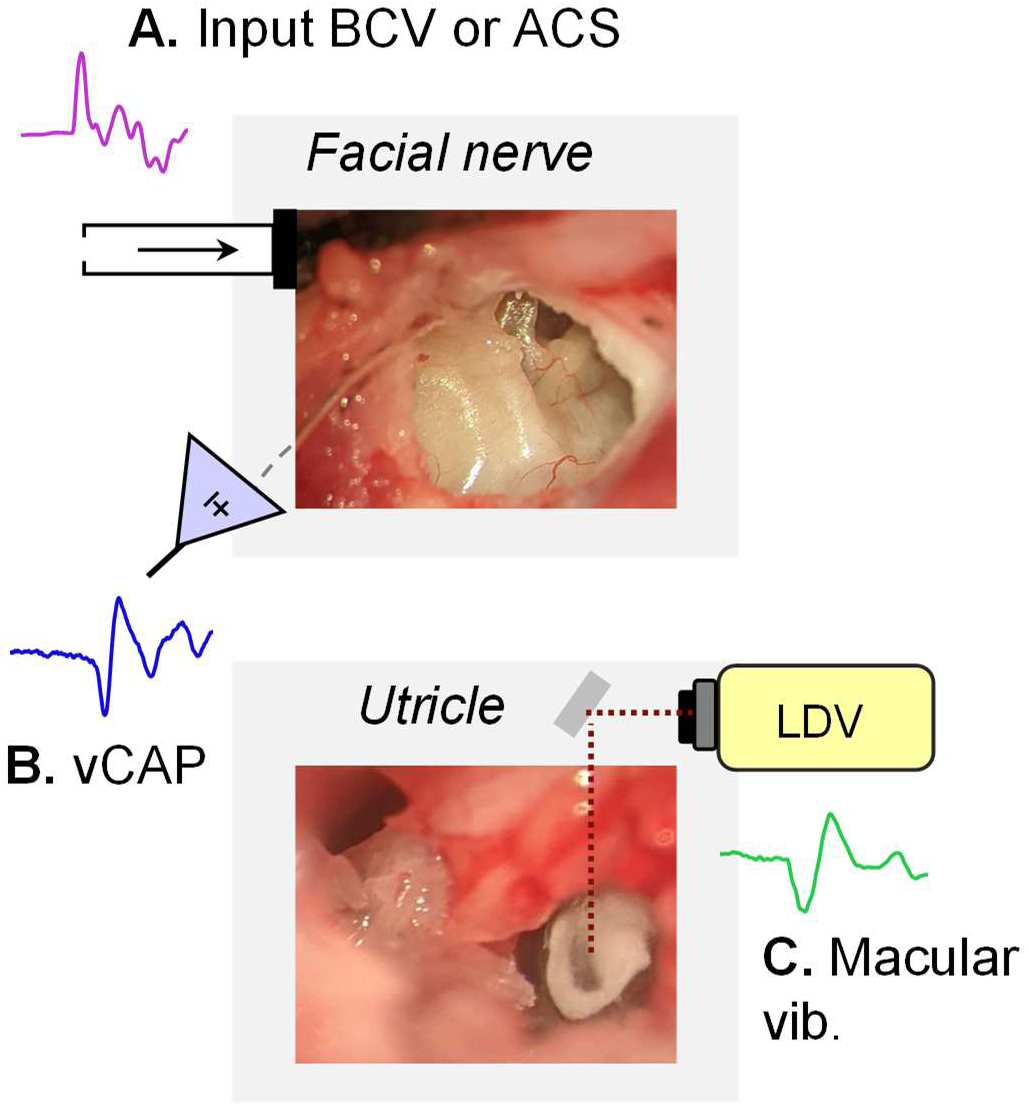
The experimental approach to record vestibular afferent and vibration responses. A. Transient BCV and ACS (magenta) were used to evoke B. synchronized vCAPs (blue) recorded from the facial nerve canal in anesthetized guinea pigs. C. Simultaneous measurements of utricular macular vibration (green) were measured via Laser Doppler Vibrometry (LDV).

### vCAP recording

To record the vestibular Compound Action Potential (vCAP), the dorsolateral bulla was exposed and opened via a postauricular surgical approach, with the guinea pig laying supine, mounted in custom-made ear-bars (modular setup using components from Thorlabs, NJ, USA). A single channel two-electrode differential recording montage was used to measure vCAPs. Here, the non-inverting (active) electrode was a fabricated 200 μm Ag/AgCl electrode that was inserted ~3mm into the bony facial nerve canal, near the vestibular branch of cranial nerve VIII (see. Fig. 1A). The inverting (reference) electrode was a custom-made Ag/AgCl electrode that was inserted into nearby neck musculature. All biopotentials were grounded via a low-resistance earth electrode placed in the nape of the neck, covered in saline-soaked gauze. vCAPs were evoked by transient pulses or tone-burst stimuli; neural origins were confirmed with chemical ablation of vCAPs following tetrodotoxin (TTX; 100μM in artificial perilymph; Sigma Aldrich, AUS) (Fig. 2).

**Fig. 2.**
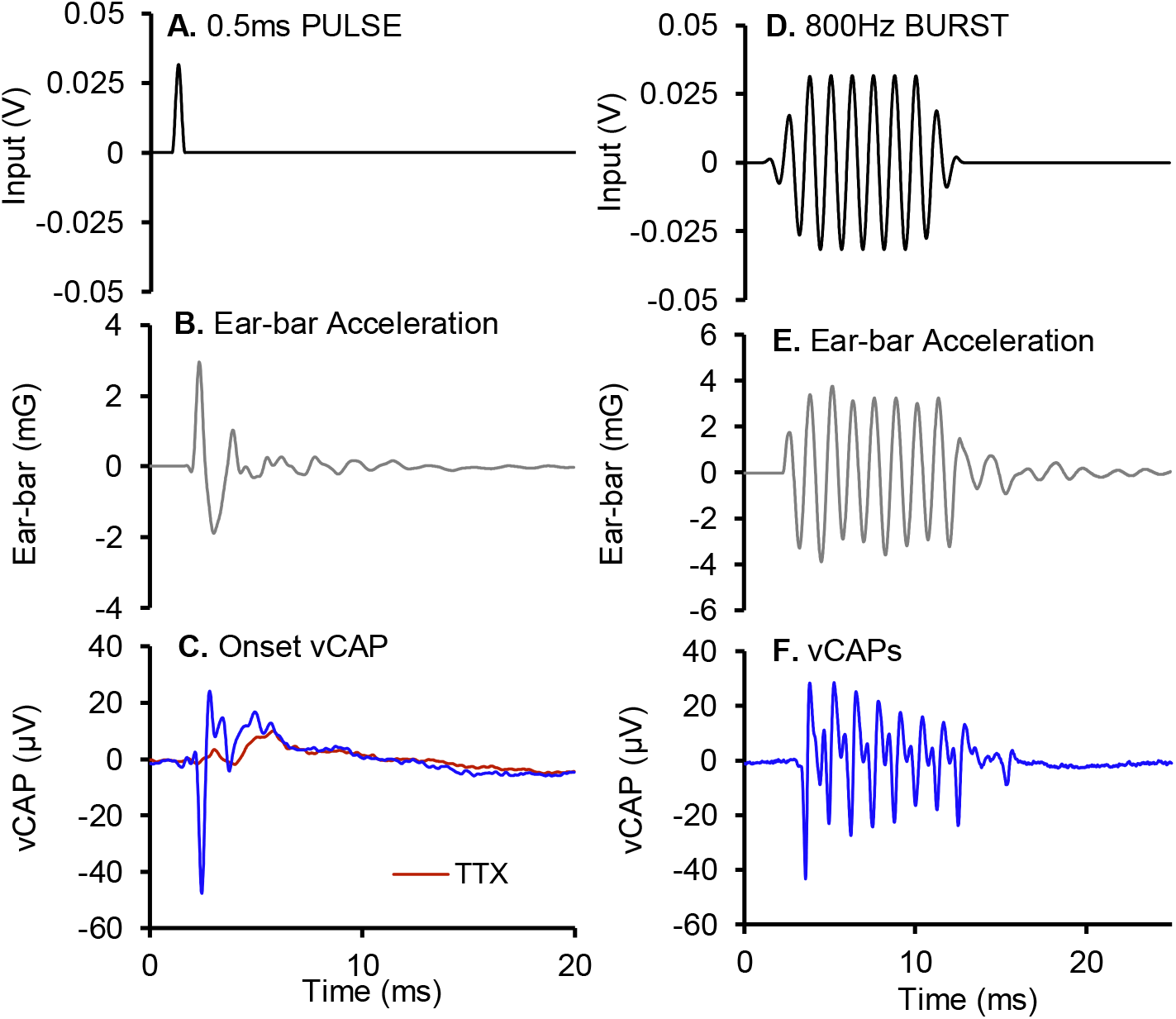
Vestibular striolar primary afferents fire synchronized action potentials at short latencies and precise phase angles relative to input BCV or ACS pulses or train bursts, respectively. A. 0.5ms BCV pulses (0.25ms rise-fall) generate punctate B. ear-bar acceleration transients which evoke robust C. vestibular nerve Compound Action Potentials (vCAPs, averaged; 100 presentations). D. 800Hz sinusoidal BCV results in a E. high-frequency acceleration burst, and a F. volley of extracellular vCAPs, also referred to as the Vestibular Nerve Neurophonic.

### VM recording

To record localized Vestibular Microphonic (VM) potentials from the basal surface of the utricular macula, the cochlea was surgically exposed and ablated using a ventral surgical approach, to provide a full view of the utricular macular epithelium under the observation of the operating microscope (see. Fig. 1, 3, & Pastras et al., 2021). The VM was measured using a two-electrode single-ended recording montage. The active electrode was an Ag/AgCl electrode placed into a pulled Borosilicate pipette with a tip diameter of ~10μm and backfilled with 250mM of NaCl. The pipette was positioned in the vestibule using a manual 3-axis micromanipulator fixed to an isolation table. The pipette electrode was guided down to the surface of the macula until touching the thin layer of perilymph above the epithelium (see. Pastras et al., 2017).

**Fig. 3:**
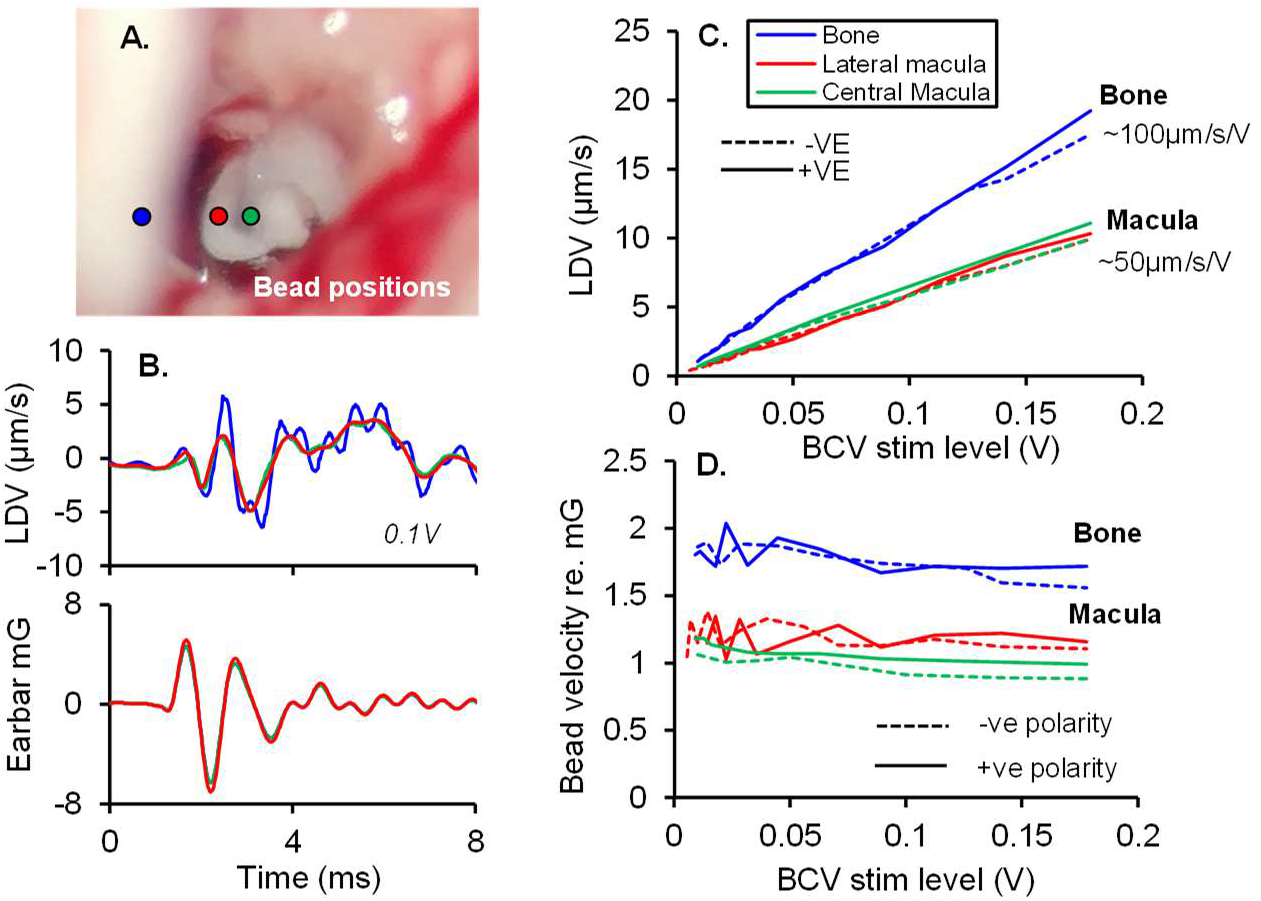
Dynamic response of the macula vs. bone to pulsatile vibration. A. Reflective microbeads were placed on the central (*green*) and lateral (*red*) macular region, and bone within the vestibule (*blue*), for LDV and accelerometer measurements during BCV. B. Representative LDV and accelerometer waveforms corresponding to 0.1V input drive to the minishaker. C. LDV Input-Output functions across BCV input drives corresponding to central and lateral macula and bone positions, for negative (dashed lines) and positive (solid lines) polarity stim. D. LDV response sensitivity relative to earbar acceleration in units of μm/s/mG, across input drives associated with different bead positions.

### Laser Doppler Vibrometry measurements

A single-point LDV (type 8338 - Brüel & Kjær, Denmark) was used to measure the dynamic response of the utricular macula and stapes during transient vibration and sound stimulation. To increase the LDV signal strength reflective microbeads (Cospheric, CA, USA) were positioned on the macula and stapes under guidance of a surgical microscope. The LDV laser beam was then directed onto the microbead targets via an adjustable optical mirror in 3D (Thorlabs, NJ, USA) (Fig. 1B). Perilymph build-up over the bead was controlled by the placement of tissue wicks into the vestibule, which minimised artifacts in the LDV recordings due to fluid surface motion effects. When recording vCAP responses, attempts were made to position the bead at the dark band at the centre of the macula, which corresponds approximately to the striolar region (Fig. 1 & 3). However, measures of macular vibration at the lateral striolar region revealed minimal to no differences to that of the central ‘striolar’ zone for pulsatile vibration (Fig. 3). This suggested that discrepancies in bead placement across animals did not alter mechanical results based on spatial tuning of the macula.

### Earbar acceleration and jerk

A triaxial piezoelectric accelerometer (Model 832M1-0200, TE connectivity, NSW, Australia) with a frequency response of 2-6000Hz and range of ±25g, was mounted to the ear-bar frame using a screw thread adapter, in the same plane as the bone-conductor (inter-aural axis). Ear-bar jerk was simultaneously calculated by taking the first derivative of ear-bar acceleration.

### Stimuli and recordings

Stimuli and responses were generated and recorded using custom-developed LabVIEW programs (National Instruments, TX, USA). BCV and ACS were generated using an external soundcard (SoundblasterX7; Creative Inc., Singapore). Analogue responses were amplified by 80dB (x10,000), with a 0.1Hz to 10kHz band-pass filter (IsoDAM8, WPI, Florida, USA) before being digitized at a rate of 40,000Hz. All responses were averaged using 100 stimulus presentations.

## Results

### vCAP sensitivity with changes in rise-time

Primary striolar afferents and their myogenic counterpart, the VEMP, have been shown to be sensitive to the very onset of the stimulus envelope and are attenuated with increases in the stimulus rise-fall time ^46^. However, the associated mechanical activation during vestibular afferent response generation under these conditions is unknown. To examine the stimulation sensitivity of the vestibular striolar afferents, vCAPs were monitored with simultaneous measures of macular epithelial vibration during changes in input drive duration (or rise-time) across several paradigms: Iso-drive, iso-macular velocity, iso-earbar acceleration, and iso-earbar jerk. The general approach was to examine the stimulation induced changes in the vCAP and associated mechanics in relation to the changes in various stimulus parameters.

### Iso-drive

Command voltages (drive) supplied to the Bruel & Kjaer minishaker as a 4ms square wave pulse were kept constant, whilst varying the stimulus rise-time between 0-2ms (Fig. 4A). vCAPs, macular vibration, earbar acceleration, and its derivative, earbar jerk, were simultaneously measured. All responses declined as a function of drive rise-time, albeit at different rates (Fig. 4B-E). Normalizing data by the shortest rise-time result revealed changes in vCAP sensitivity (Fig. 5, red) were closely correlated with the changes in macular velocity (Fig. 5, blue circles). Both the vCAP amplitude and macular velocity declined approximately linearly with increases in drive rise-time for all stimulus intensities tested (Fig. 5A & B). By contrast, earbar acceleration and earbar jerk declined nonlinearly with increased rise-time, with earbar jerk displaying a greater rate of decline than acceleration, especially for brief rise-times <1ms (Fig. 5A & B). Response amplitudes normalized to the vCAP (normalized by the vCAP magnitude in A-B) reveal a close correlation between vCAP and macular velocity magnitudes (Fig. 5C & D). Of the two earbar kinematic variables, the vCAP (and macular velocity) scaled most closely with earbar acceleration (magenta) compared to earbar jerk (grey), for all stimuli used during the iso-input drive paradigm (Fig. 5C & D).

**Fig. 4.**
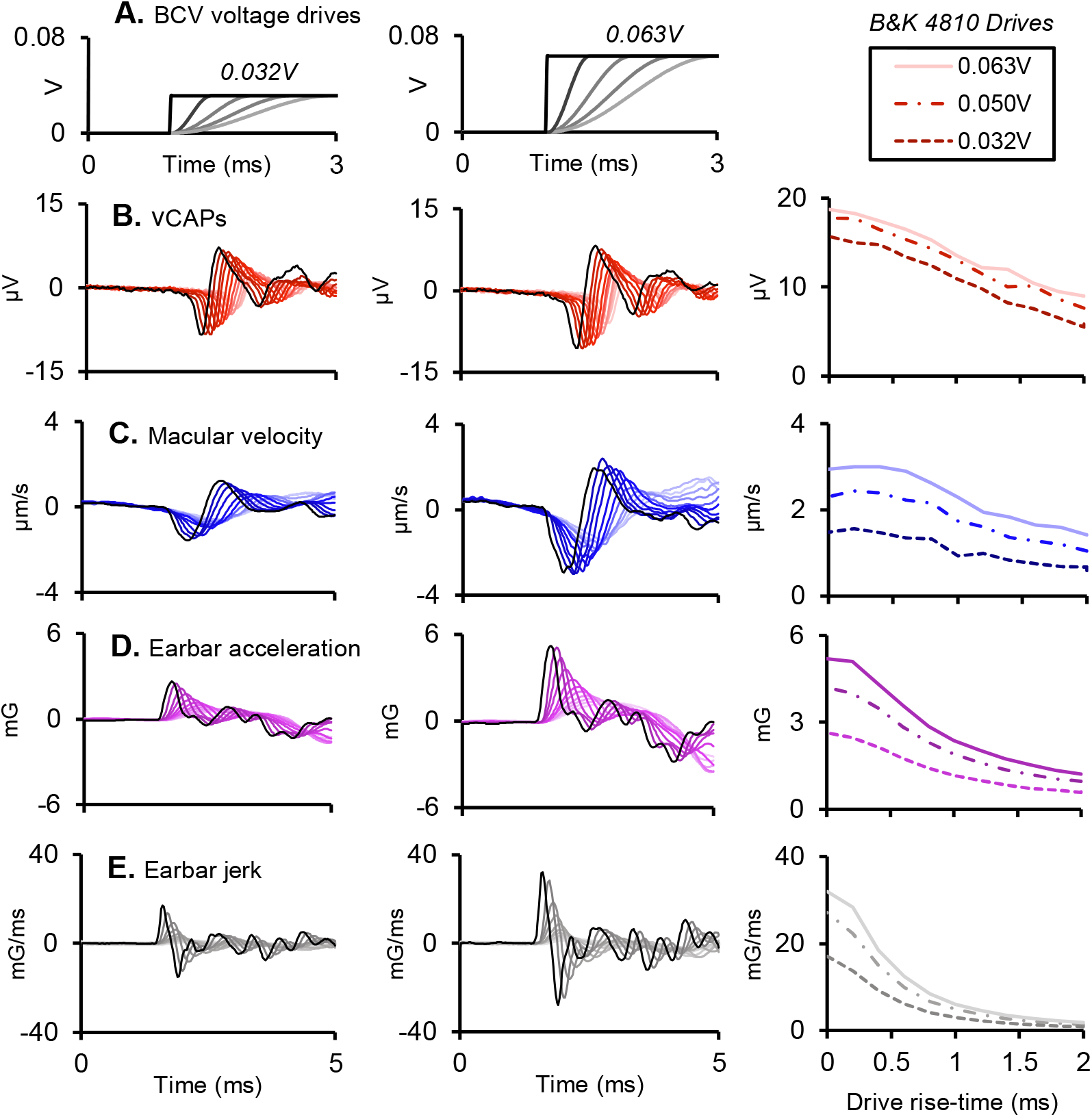
Macular responses during BCV iso-drive at different stimulus intensities, between 0.032V-0.063V. A. The BCV voltage drive to the mini shaker was kept constant (iso-drive) whilst the stimulus rise fall-time was varied (0-2ms; 0-50%) for a 4ms BCV pulse. Simultaneously measured B. vestibular compound action potentials (red), C. macular velocity (blue), D. earbar acceleration (magenta), and its derivative, E. earbar jerk (grey). Responses in the left and middle panel correspond to the lowest (0.03V) and highest (0.06V) stimulus intensity, respectively.

**Fig. 5.**
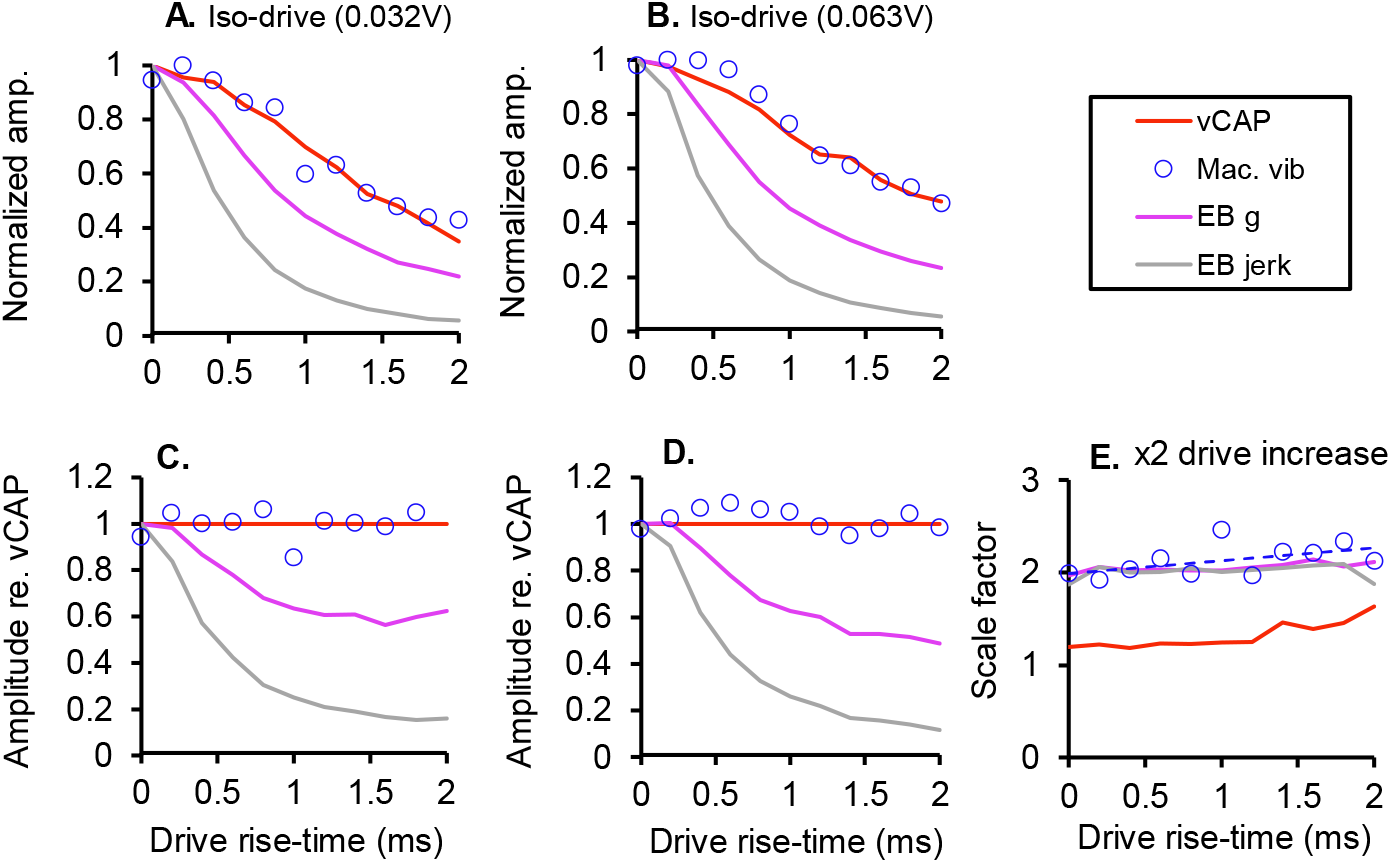
Normalized data from Fig. 4 showing macular response scaling and stimulation sensitivity. A, B. Responses during iso-drive stimulation of the macula normalized to maximal amplitude for 0.032V and 0.063V BCV stimulus intensities, respectively. C, D. Response amplitudes normalized to vCAP amplitude, corresponding to data in A,B, respectively. E. Response scaling with changes in stimulus rise-time associated with a x2 BCV drive increase.

Doubling the BCV input command voltage drive (0.03V vs 0.06V) resulted in a doubling of the mechanical response sensitivity, which included macular velocity, earbar acceleration, and earbar jerk (Fig. 5E). By comparison, the same two-fold increase in BCV drive resulted in a compressive scaling (scale factor) of the vCAP (~1.2-1.5x increase), suggesting vestibular neural output is nonlinear, whereas macular mechanics is linear and passive.

### Iso-macular velocity

Macular response sensitivity was further characterized by monitoring vCAP response amplitudes during an iso-macular velocity paradigm with associated changes in input drive rise-time. Here, the voltage drive to the minishaker was varied to produce a fixed first negative macular velocity peak (N1) associated with changes in stimulus rise-time between 0 and 2ms (Fig. 6A). Macular displacement and earbar velocity increased linearly as a function of increased BCV drive rise-time associated with constant macular velocity (Fig. 6B & E). vCAP peak-peak amplitude scaled closely with onset macular velocity and remained consistent between rise-times of 0 to 1ms. However, vCAP sensitivity began to decline between 1 and 2ms, associated with longer BCV rise times (Fig. 6C). Earbar acceleration declined linearly between 0 and 2ms (Fig. 6D), and earbar jerk, declined exponentially (Fig. 6F) over a 2ms change in stimulus drive rise-time.

**Fig. 6.**
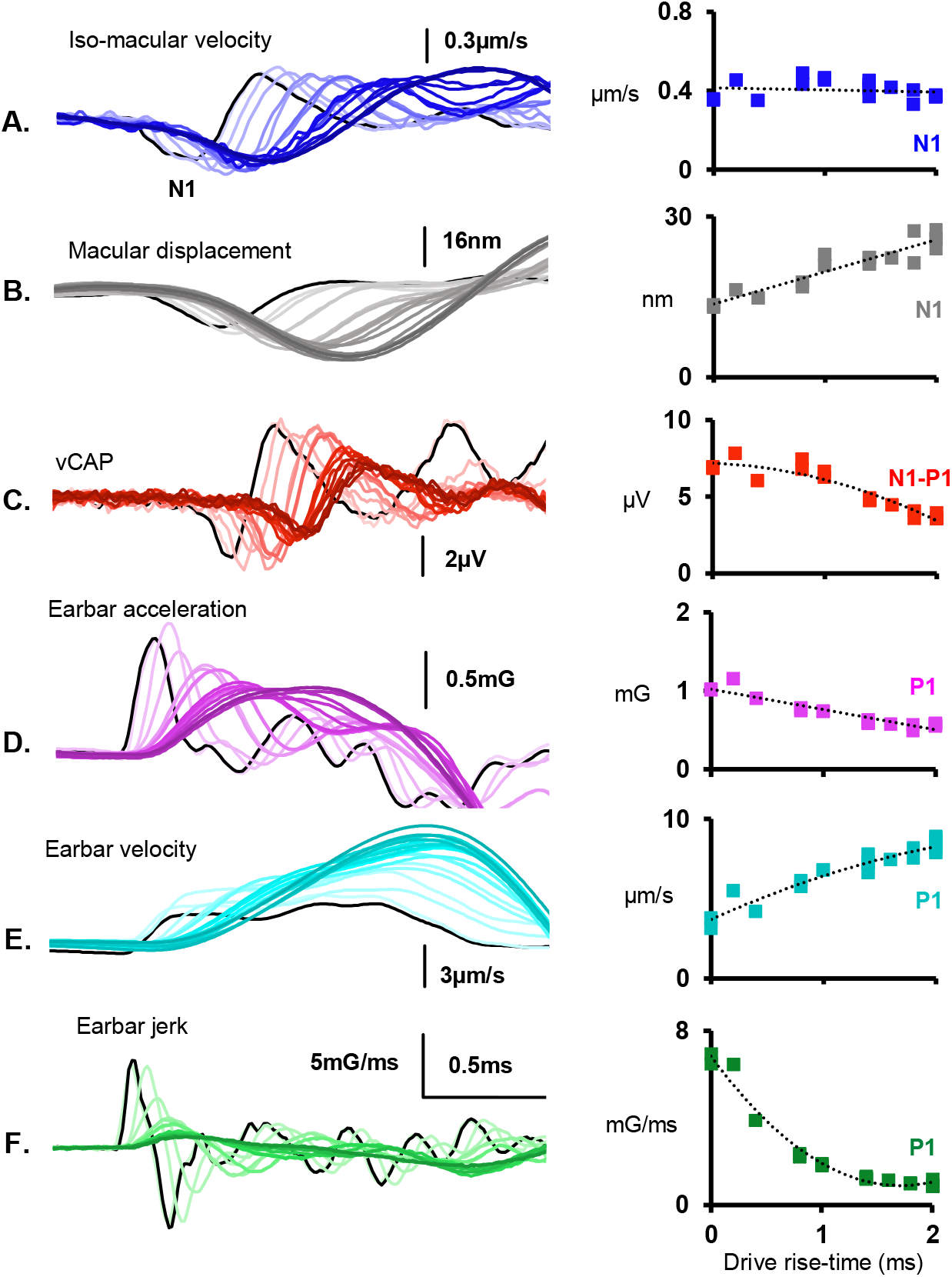
Iso-macular velocity and associated response waveforms and magnitudes. A. The magnitude of macular velocity (blue) was kept constant (*iso-macular velocity*) with changes in drive rise-time (0-50%; 0-2ms) associated with a 4ms BCV pulse. B. Its integral, macular displacement (grey) was quantified, along with C. synchronized vCAPs (red) recorded from the facial nerve canal, D. earbar acceleration (magneta), E. its integral, earbar velocity (cyan), and F. earbar jerk (green).

### Iso-earbar acceleration and jerk

To further probe the kinematic sensitivity of the vCAP with regards to the earbar (and cranium), vCAPs were recorded while keeping earbar acceleration (Fig. 7), or earbar jerk (Fig. 8), constant as rise-time was varied. For a fixed earbar acceleration (1mG, Fig. 6A), earbar jerk declined exponentially (Fig. 7C), while earbar velocity and macular displacement increased sigmoidally, beginning to saturate at long duration drive rise-times (Fig. 7B & F). vCAPs increased in amplitude with brief drive rise-times between 0 and 0.5ms, but began to decline with longer rise-times, between 0.5 and 2ms (Fig. 7D). This is consistent with a switch in vCAP sensitivity from temporal bone acceleration to jerk for longer BCV durations. Macular velocity increased as a function of input rise-time, up until 1.5ms, where epithelial vibration began to decline (Fig. 7E), consistent with the saturation of macular displacement at long rise times (Fig. 7F). For a fixed earbar jerk (6mG/ms, Fig. 8A), vCAPs scaled approximately with earbar acceleration and macular velocity for brief rise-times (0-0.5ms) (Fig. 8B, C & D). However, for longer duration rise-times (>0.5ms), vCAP amplitudes saturated and scaled with earbar jerk (Fig. 8A-B). A fixed earbar jerk resulted in a proportional scaling of earbar velocity with macular displacement (Fig. 8D & F), and a similar scaling relationship of earbar acceleration to macular velocity (Fig. 8C & E).

**Fig. 7.**
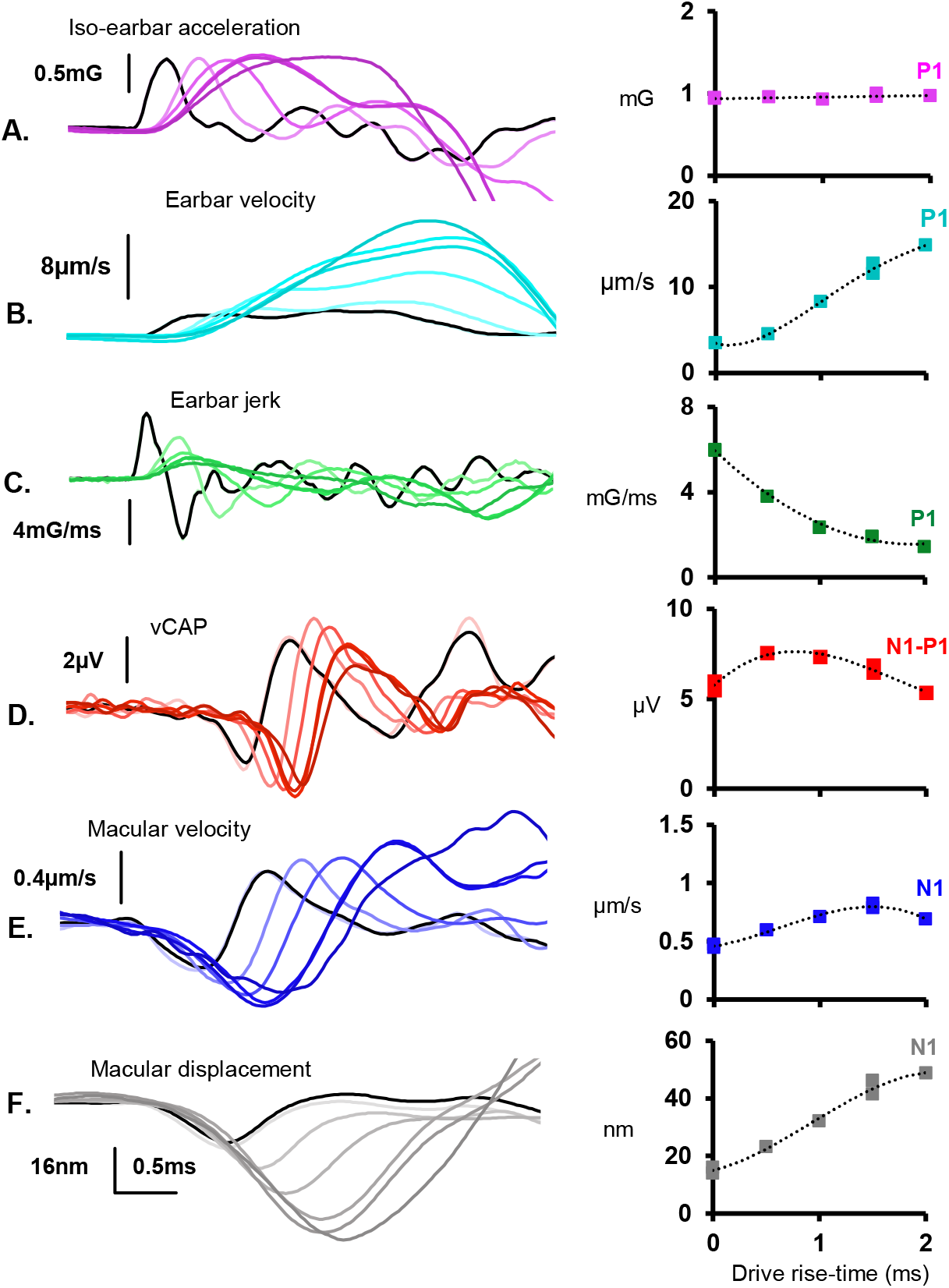
Iso-earbar acceleration and associated response waveforms and magnitudes. A. The magnitude of earbar acceleration (magenta) was kept constant (*iso-earbar acceleration*) with changes in input drive rise fall time (0-50%; 0-2ms) associated with a 4ms BCV pulse. B. Its integral, earbar velocity (cyan), and C. its derivative, earbar jerk (green) was quantified, along with D. vCAPs (red), E. macular velocity (blue), and F. its integral, macular displacement (grey).

**Fig. 8.**
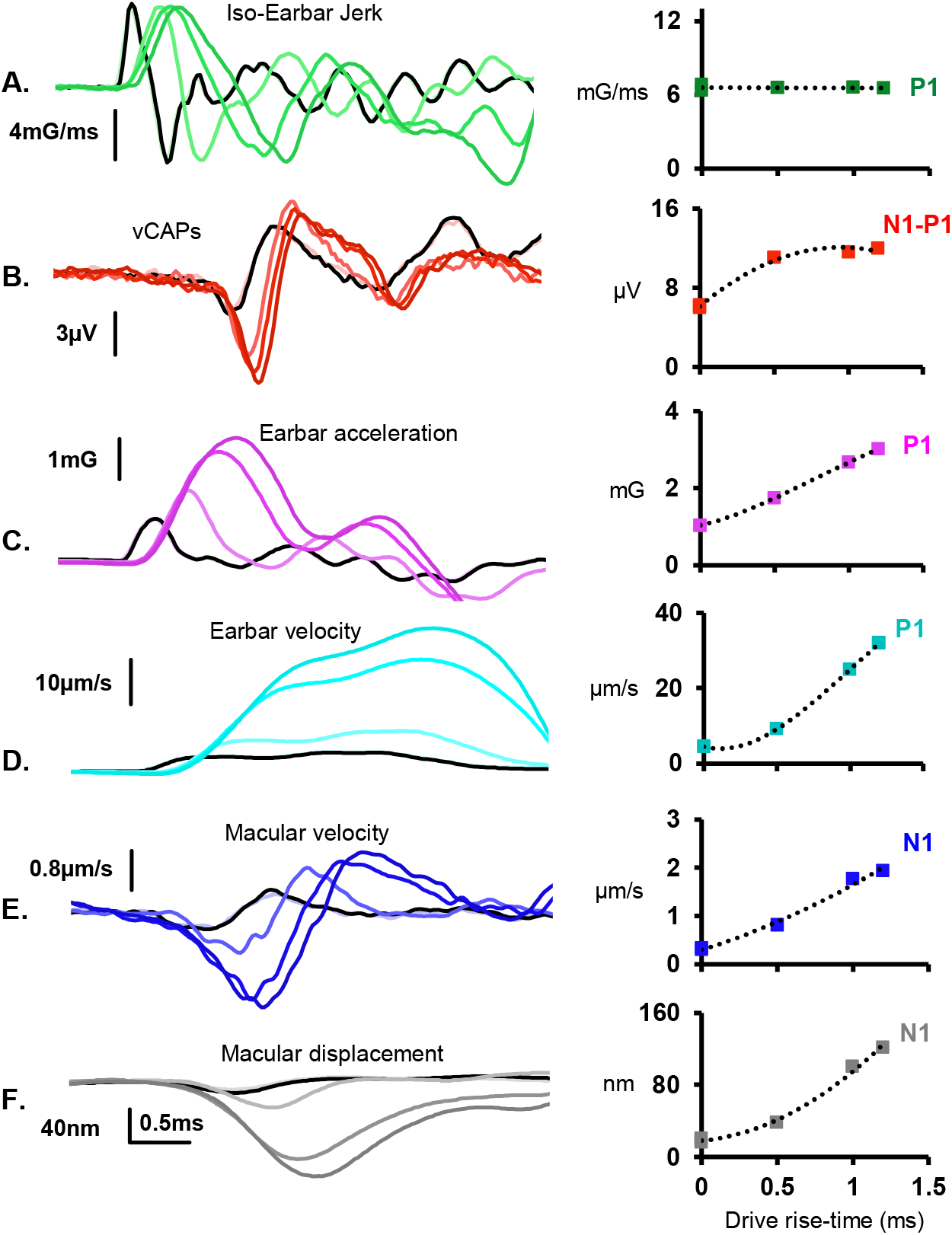
Iso-earbar jerk and associated response waveforms and magnitudes. A. The magnitude of earbar jerk (green) was kept constant (iso-earbar jerk) with changes in input drive rise fall time (0-2ms) associated with a 4ms BCV pulse. B. vCAPs were measured from the facial nerve canal, (red), alongside C. earbar acceleration (magenta), D. earbar velocity (cyan), E. macular velocity (blue), and F. its integral, macular displacement (grey).

Waveform and magnitude data from Figs 6-8 were normalized for each paradigm (Fig. 9), showing the relative scaling of each response as a function of input drive rise-time. For iso-macular velocity, iso-earbar acceleration and iso-earbar jerk (Fig. 9A-C), the vCAP scales almost in proportion to macular velocity and earbar acceleration for short duration rise-times (<1ms). However, for longer duration rise-times, the scaling of the vCAP becomes divergent with macular velocity (and earbar acceleration) and begins to approximate the scaling of earbar jerk.

**Fig. 9:**
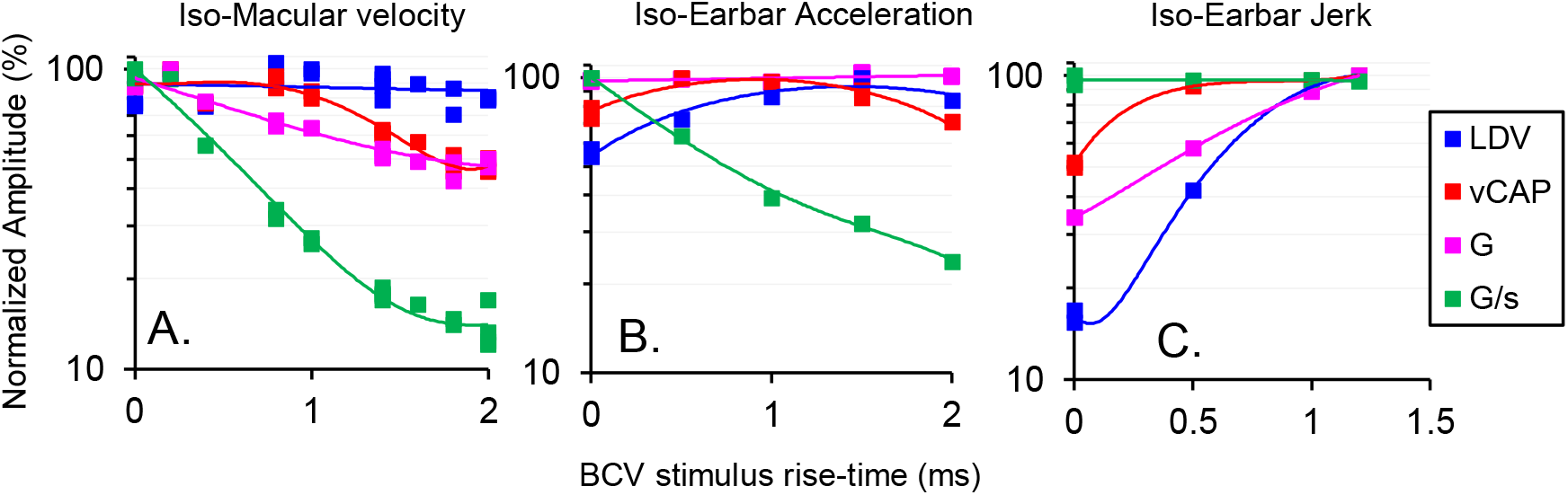
Normalized amplitudes associated with changes in stimulus paradigm. Macular velocity (blue), vCAP (red), earbar acceleration (magenta), and earbar jerk (green) for A. iso-macular velocity, B. iso-earbar acceleration, and C. iso-earbar jerk. Normalized amplitude plots in A-C correspond to waveforms in Figs 6-8, respectively.

BCV chirps were used to assess the relationship between mechanical activation of the macula and vCAP generation for more complex vibrational stimuli. A 10ms backward chirp (0ms rise-time) with broadband spectra (Fig. 10A,B) generates a highly synchronous vCAP with robust macular vibration, and relatively broadband earbar vibration (Fig. 10C-F). Increasing the chirp stimulus rise-time from 0ms to 5ms completely abolishes the vCAP response, leaving behind a contralateral Auditory Brainstem Response (ABR), which disappears following contralateral cochlear ablation (data not shown). The ABR response scales closely, in timing and amplitude, with the mid-latency (high frequency) components of earbar acceleration and jerk (Fig. 10F), whereas the vCAP scales closely with onset macular velocity and onset earbar acceleration. These results reveal that transient onset stimuli are needed to produce sufficient macular vibration for the synchronization of otolithic afferent responses and the generation of sensory vCAPs.

**Fig. 10:**
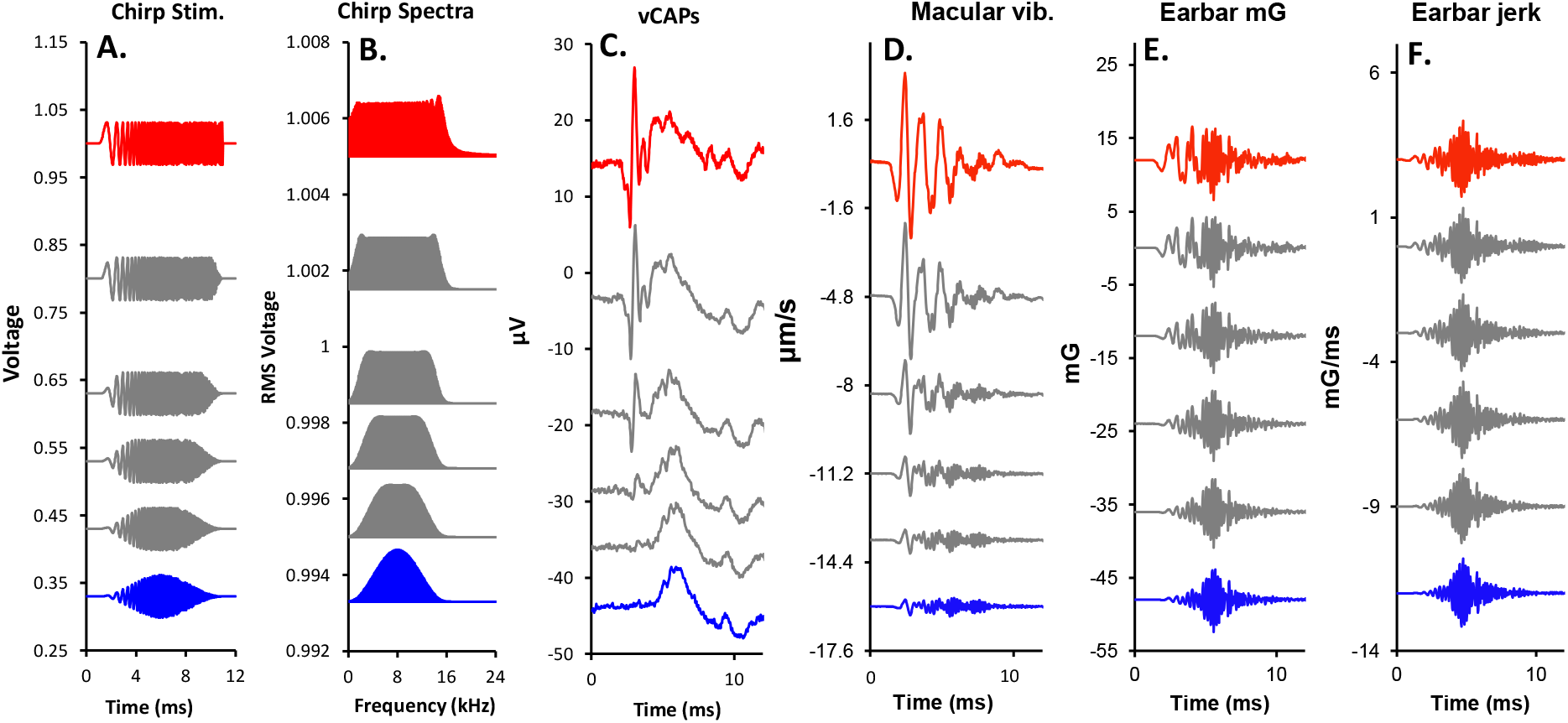
vCAP sensitivity to broadband chirps. A. 10ms BCV chirps with varying rise-times, B. Chirp power spectra, C. measured vCAPs, D. macular velocity using LDV, E. earbar acceleration, and F. jerk.

To further probe the relevant stimulus characteristics for evoking transient vestibular responses using broadband input, both backward and forward chirps were used to evoke vCAPs, with corresponding measures of skull vibration (Fig. 11A-D). Data reveal that the latency and generation of the vCAP closely follows the timing of the low frequency component of the broadband stimulus, with most of its spectral power falling below 1kHz (Fig. 11A). At its simplest, this follows from the undamped low-pass biomechanics of the otoliths, with a natural frequency around 500Hz.

**Fig. 11:**
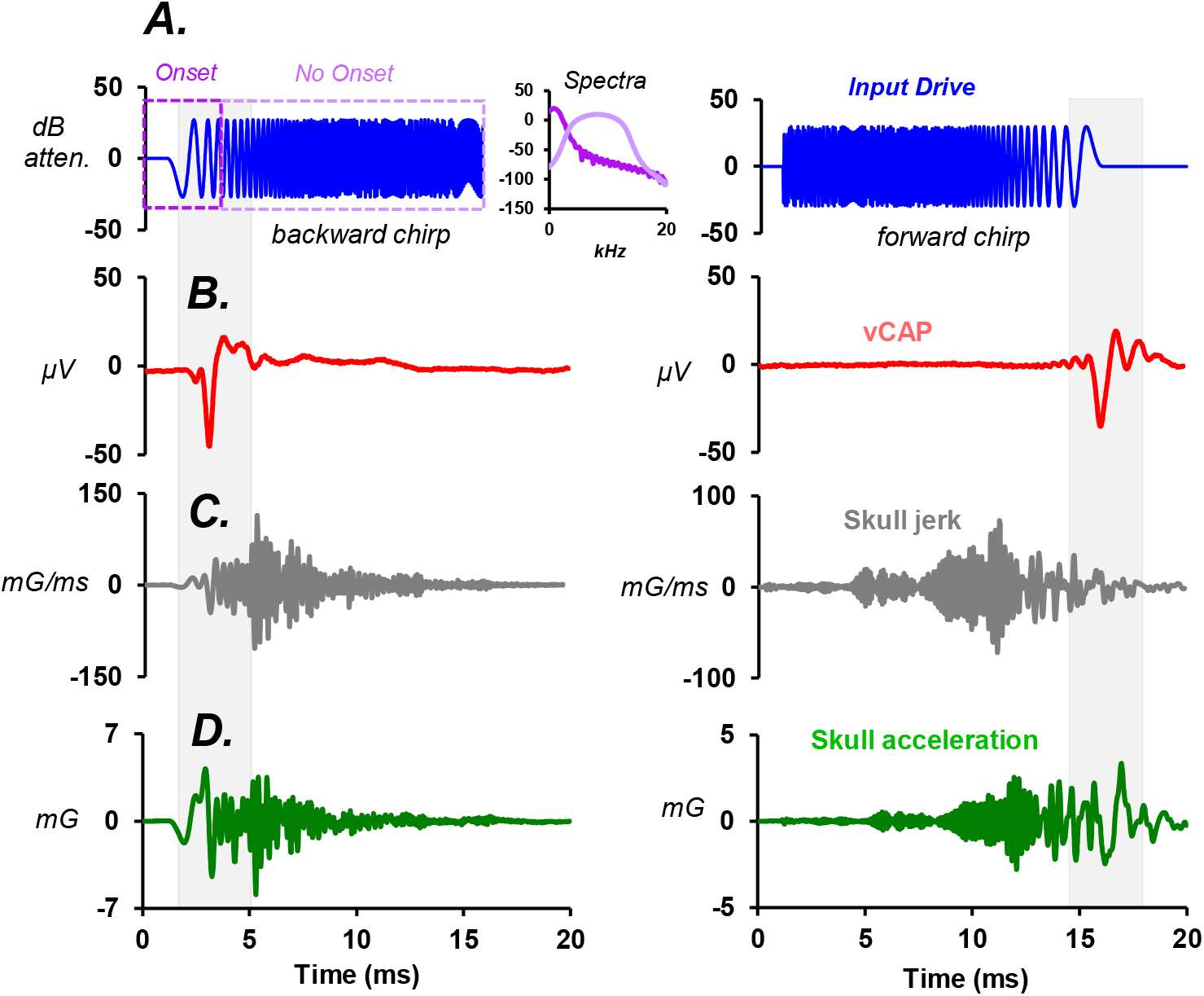
Chirp direction and vCAP generation. A. Backward and forward BCV chirps generated B. vCAPs which followed the low-frequency component of the broadband stimulus. C. Simultaneously measured skull jerk, and D. acceleration. Inset: Chirp stimulus power spectrum (hanning window).

### BCV vs ACS vCAP IO functions

To characterize the sensitivity of irregular striolar afferents to increasing levels of mechanical stimulation, pulsatile BCV and ACS stimuli (0.5ms duration, 0.25ms rise-fall) were used to evoke vCAPs, with simultaneous measurements of macular epithelial vibration using LDV. Representative waveforms from one example animal show typical BCV (Fig. 12A-B) and ACS (Fig. 12D-E) evoked vCAP responses, along with macular velocity measurements to a range of stimulus intensities. The corresponding IO function (vCAP amplitude vs macular velocity) of these example plots is shown below (Black lines; Fig. 12C & F), along with the IO function from 18 (BCV) and 11 (ACS) other animals. It should be noted that in this intra-animal comparison, vCAP responses are approximately 5x larger for BCV than ACS stimuli, for a similar level of macular velocity. Furthermore, when comparing vCAP IOs across animals and stimuli (Inter-animal; Fig. 12C, F & Inset), the overall sensitivity (slope) of vCAP IOs was greater for BCV than ACS stimuli (Fig. 12C & F), which is represented in the Figure 12 inset, with ACS & BCV overlaid.

**Fig. 12.**
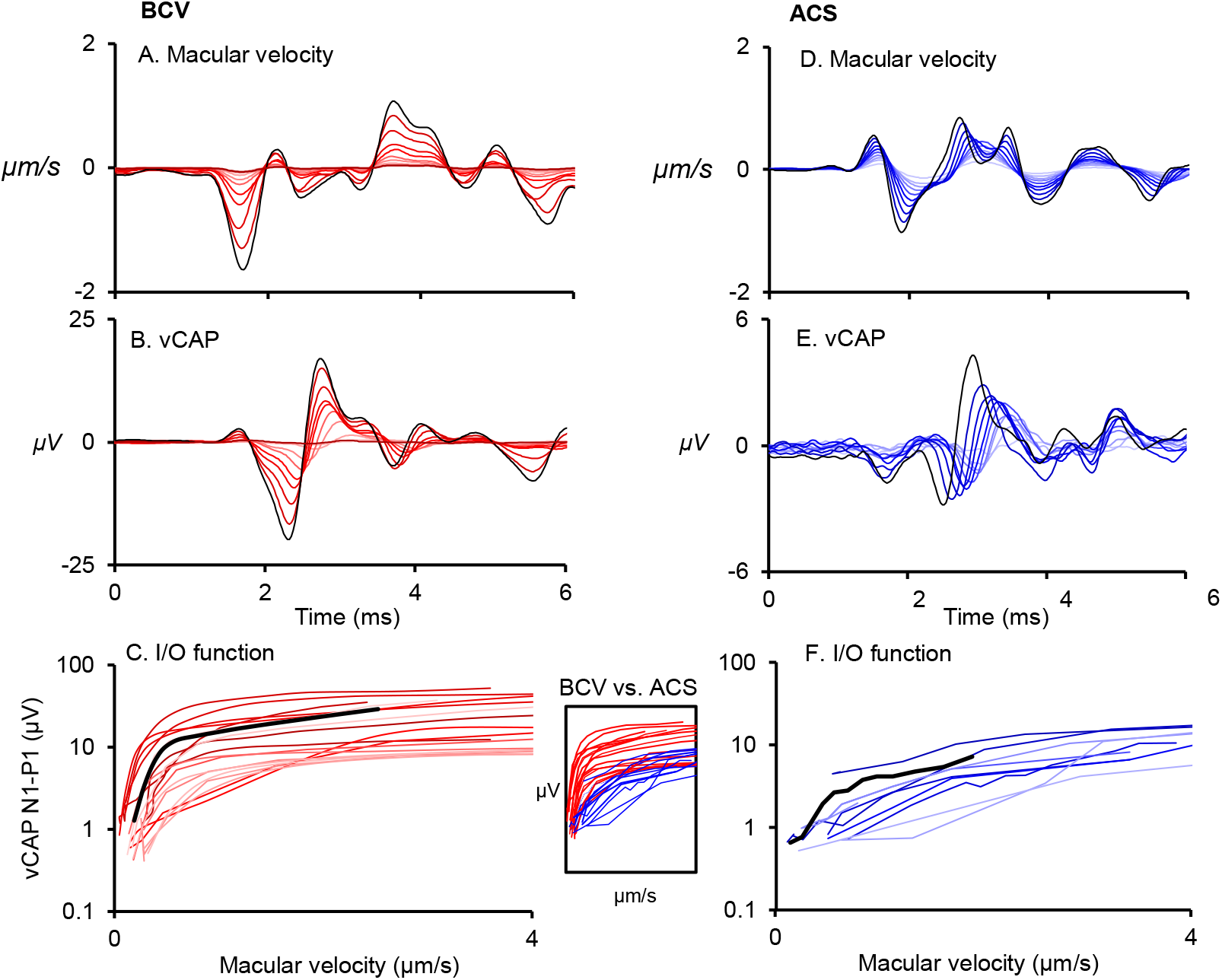
Inter-animal comparisons of vCAP Input-Output (IO) functions for BCV and ACS. A. Representative BCV macular velocity and B. vCAP waveforms in 1 animal, with the associated IO curve displayed as the black trace below. C. BCV IO functions across 18 animals. D. Representative ACS macular velocity and E. vCAP waveforms in the same animal, with its corresponding IO curve displayed as the black trace below. F. ACS IO functions across 11 animals. Inset: Overlay comparison of BCV and ACS IO functions. Although, IO thresholds are similar, IO slopes are larger for BCV data.

Given, ACS relies on impedance matching between the atmosphere and the viscous environment of the inner ear, we wanted to investigate whether sensitivity differences between BCV and ACS vCAPs where due to reduced fluid coupling between the stapes footplate and the macula, providing inadequate mechanical drive to the vestibular hair cells.

To investigate these effects, fluid was removed from within the vestibule for both BCV and ACS responses (Fig. 13A & B) and was compared to recordings made when perilymph was fully covering the macular surface (Fig. 13C), in the same animal. With fluid in the vestibule, direct LDV measures of macular vibration were infeasible. Instead, as a proxy, we measured stapes footplate vibration with LDV (Fig. 13C). Without fluid coupling, the vCAP amplitude and IO slope were much smaller for ACS than for the BCV stimulation (Fig. 13D). When fluid coupling was improved, the ACS vCAP amplitude and IO slope increased to the level of the BCV vCAP IO (Fig. 13D & E) revealing the differences in the ACS vCAP were likely due to altered fluid coupling and changes impedance matching. This reveals the sensitivity of the utricular nerve in generating synchronized action potentials of short latency is equivalent for punctate ACS and BCV stimuli in the guinea pig. These recordings also provide good estimates for the level of mechanical vibration of the macula and stapes at the threshold of action potential generation in the anaesthetized guinea pig. Results reveal that the magnitude of macular velocity for vCAP threshold is ~0.3μm/s for BCV and ACS, and orders of magnitude greater for stapes velocity (~25μm/s) associated with ACS (Fig. 13D).

**Fig. 13.**
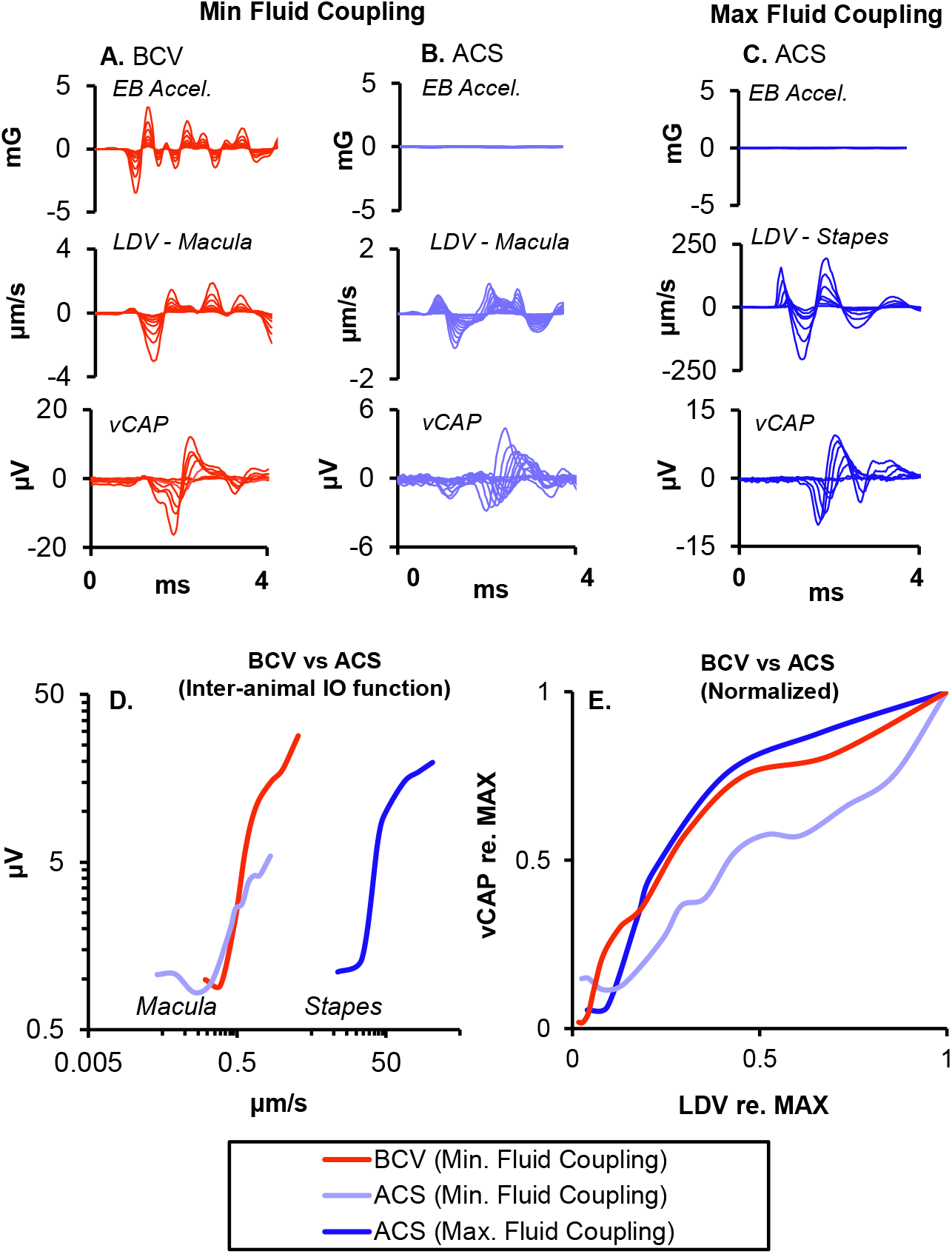
A. Intra-animal comparisons of vCAP Input-Output (IO) functions for BCV and ACS for Minimal and Maximal Fluid Coupling between the stapes and macula. A, B. Earbar acceleration, macular vibration and vCAP waveforms associated with BCV and ACS, respectively, during Minimal Fluid Coupling. C. Earbar acceleration, macular vibration and vCAP waveforms during ACS for Maximal Fluid Coupling. D. Corresponding vCAP IO functions associated with waveforms in A,B, & C, above. E. Normalized vCAP IO function data.

### vCAP sensitivity with changes in frequency

BCV and ACS tone bursts are routinely used in the neuro-otology clinic to evoke vestibular reflex responses, such as the VEMP, as a part of a standard assay of otolith function. Although there are mixed data on VEMP tuning curves, likely due to differences across recording setups, the optimal VEMP frequency is generally reported to be around 500Hz. However, the basis for this tuning is unclear relative to mechanical input and the generation of MET currents. To characterize the vCAP frequency response as a proxy from utricular afferent sensitivity across frequency, BCV tone bursts between 100-2000Hz of varying intensity levels were used to evoke a fixed amplitude vCAP, with simultaneous measures of epithelial vibration (iso-vCAP frequency tuning curve; Fig. 14A).

**Fig. 14.**
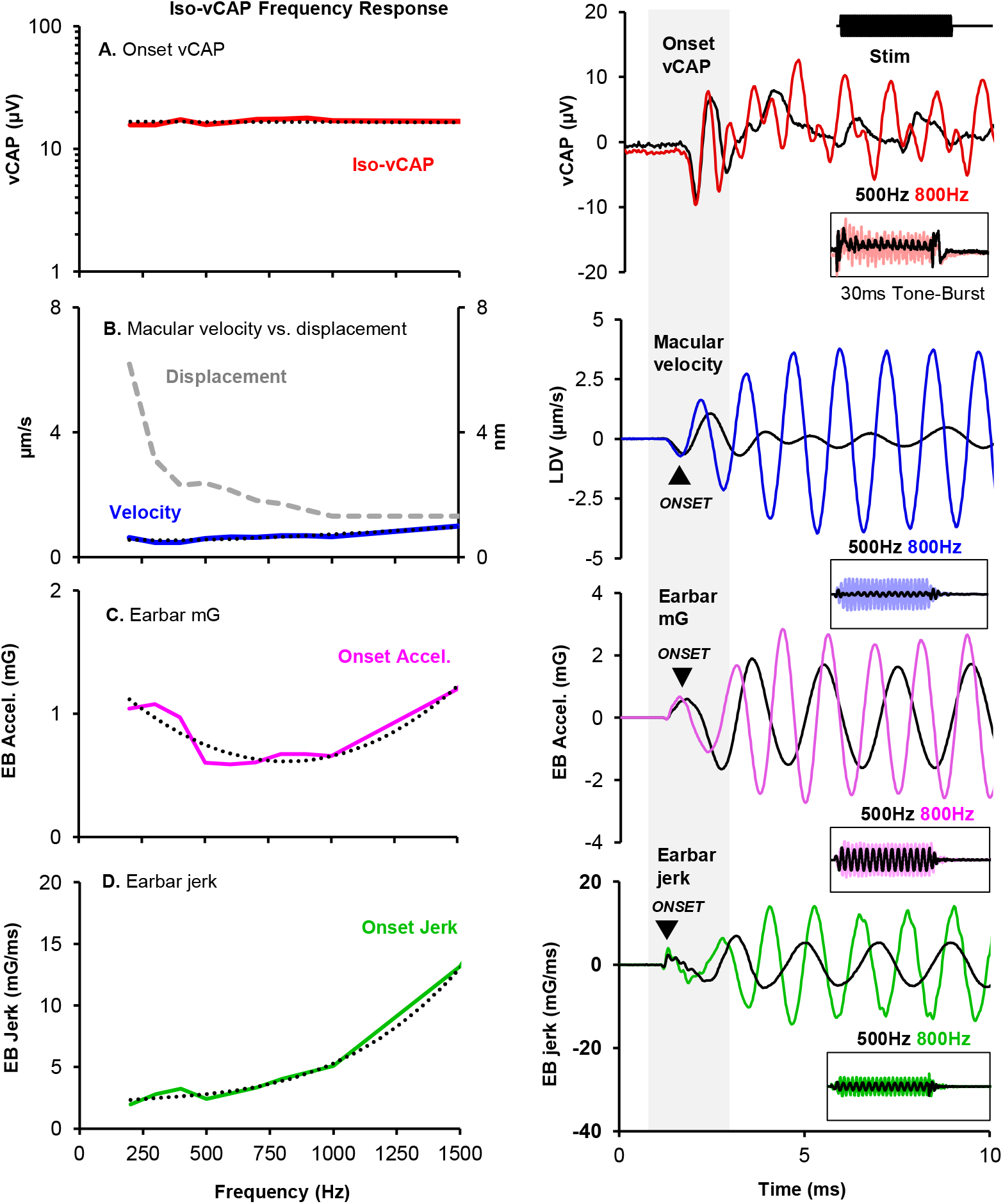
Iso-vCAP frequency response tuning curve. A. Onset vCAPs were kept constant during a 30ms BCV tone burst (0ms rf) across frequency (up to 1.5kHz), with simultaneous measurements of B. macular velocity, C. earbar acceleration, and D. its kinematic derivative, earbar jerk. E. Representative waveform comparisons for the onset vCAP, macular velocity, earbar acceleration, and earbar jerk associated with a 500Hz (black) and 800Hz (coloured) tone-burst, respectively (10ms window). Inset: Entire 50ms time-domain window of the tone-burst response.

Associated macular velocity, macular displacement, earbar acceleration, and earbar jerk were also plotted against the frequency of the BCV stimulus (Fig. 14B-D). Results reveal that for an iso-onset vCAP response (Fig. 14A), the associated onset macular velocity (taken as the initial N1 transient bump) remains relatively flat across frequency (Fig. 14A & B), suggesting that the vCAP scales with macular velocity for transient stimuli such as onset tone-bursts and pulses. By comparison, macular displacement declined exponentially with stimulus frequency, with displacement being largest at low frequencies. Earbar acceleration approximated a parabolic function over frequency (Fig. 14C), whereas earbar jerk increased exponentially (Fig. 14D). At low frequencies (<450Hz), earbar jerk was relatively flat and had comparable scaling to the onset vCAP, consistent with the finding that vestibular afferents scale with jerk for spectral power below the natural frequency of the otoliths.

### Iso-VM Frequency tuning curves

To test the extent vCAP tuning curve was related to pre-synaptic hair cell responses, Iso-macular velocity tuning curves were recorded in the same animal. Voltage drives to the mini shaker were programmatically altered to produce a constant macular velocity across frequency from 100-2000Hz (Fig. 15A), whilst simultaneously recording the VM, macular displacement, earbar acceleration, and total harmonic distortion of the recording system (Fig. 15B-D). Results reveal that for a fixed macular velocity across BCV frequency (Fig. 15A), VM amplitude and sensitivity is closely correlated with macular displacement (Fig. 15B), and this tuning is independent of temporal bone acceleration and distortion in the recording setup (Fig. 15C-D).

**Fig. 15.**
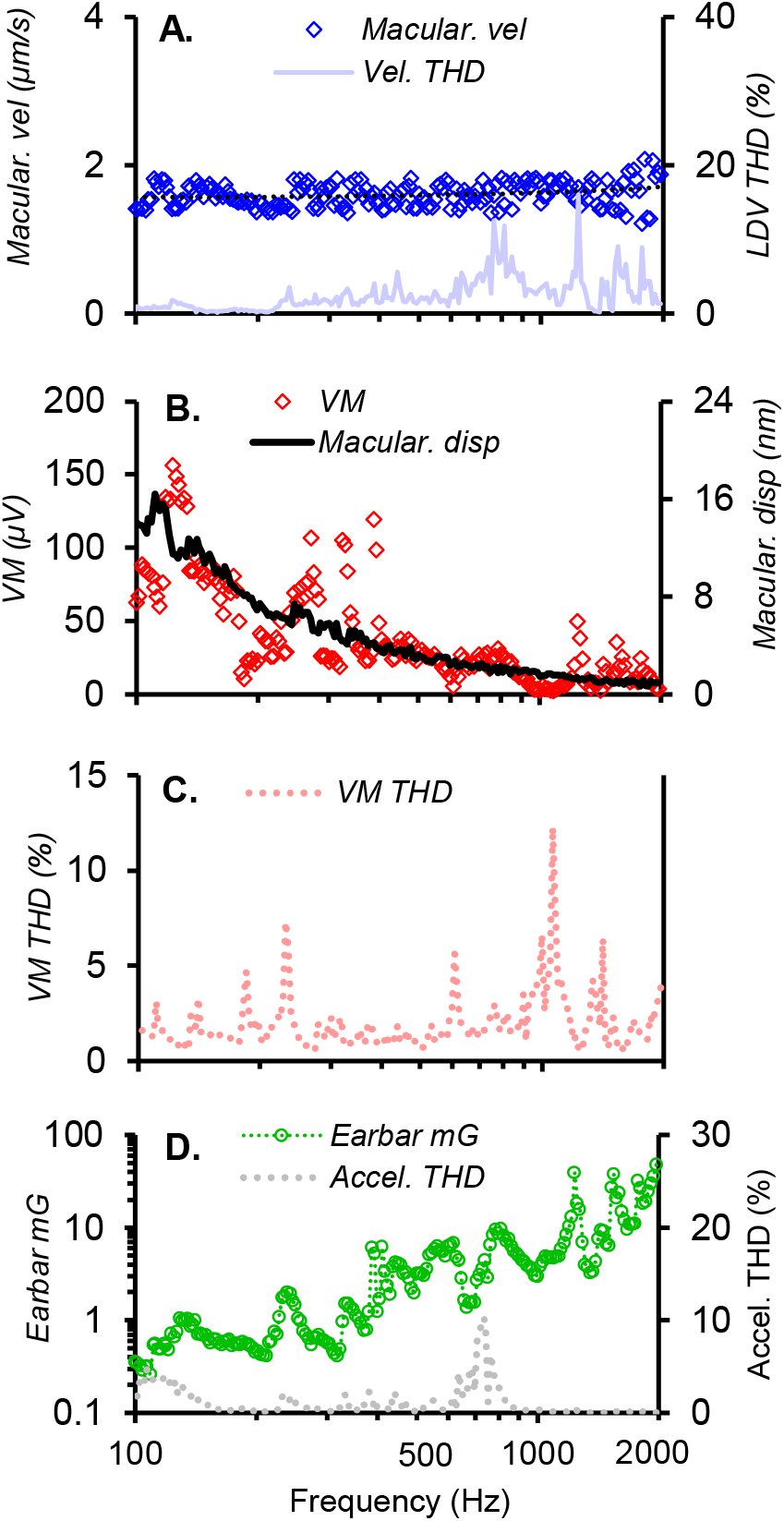
VM frequency tuning curve. A. Macular velocity (blue) was kept constant over the full bandwidth (iso-macular velocity), with simultaneous measurements of LDV total harmonic distortion (light blue), B. vestibular microphonics, associated macular displacement, C. VM THD, D. earbar acceleration and associated THD.

## Discussion

Punctate linear vibration stimuli such as brief hammer or finger taps^47^, transient BCV or ACS stimuli, and tone-bursts delivered by speakers or audiometric bone transducers are routinely used in the clinic or laboratory to evoke robust VEMP and vCAP responses. However, the mechanisms underlying these neurophysiological responses are not well understood. In the present work, we directly measured mechanical vibration of the macula, VMs and vCAPs in guinea pigs to determine how clinically relevant BCV and ACS stimuli evoke synchronized action potentials in the utricular nerve.

We first examined the relationship between the BCV stimulus and the vibration of the macula by comparing the peak macular velocity to the peak linear ear-bar acceleration (G) and jerk (G/s) for a series of stimulus strengths. Results in Figs. 6 and 7 demonstrate the peak macular velocity increases roughly in proportion to the acceleration stimulus, consistent with the prediction of simple one degree-of-freedom (1-DOF) models of the utricle for stimuli at or below the corner frequency ^48,49^. Mechanical simulations using a 2-DOF model of utricular mechanics^50^ reproduce the LDV velocities reported here, further confirming that the utricle behaves as a simple inertial sensor that responds to acceleration for short stimuli. The present LDV measurements are consistent with a slightly underdamped mechanical response, exhibiting low-pass sensitivity to sinusoidal inter-aural vibration with a corner frequency near 500 Hz ^50,51^.

In terms of the applied BCV stimulus, present results reveal the vCAP magnitude scales most closely with acceleration for short drive rise-times (<1ms), and switches to approximate linear jerk for longer duration rise-times (>1ms). These results were reproduced across three experimental paradigms, which included iso-macular velocity (Fig. 6), iso-earbar acceleration (Fig. 7), and iso-earbar jerk (Fig. 8). For short rise-times, the vCAP magnitude scaled most closely with macular velocity, and earbar acceleration, rather than other kinematic components such as macular displacement or earbar jerk (or macular acceleration; not shown, or earbar displacement; also, not shown). Hence, for very brief BCV stimuli, linear acceleration was the *adequate stimulus* to generate synchronized vCAPs in the present guinea pig experiments. However, at longer stimulus pulse widths, vCAP scaling approximated the time-derivative of earbar acceleration, which is consistent with previous VsEP experiments in rodents where linear jerk was clearly identified as the adequate stimulus to generate evoked responses ^42,52^. Despite this, there are key differences between the present report and previous VsEP studies that likely underlie the difference in sensitivity including: 1) Animal model: use of the guinea pig (*Cavia porcellus*) in the present report vs. mice (C57BL/6J) or rats (Sprague Dawley); 2) Stimulus: ~3mG inter-aural acceleration at ~20mG/ms in the present report vs. ~2000mG nasal-occipital acceleration at ~1000mG/s jerk in a supine position; 3) vCAP recording: non-inverting (active) electrode inserted in the facial nerve canal in the present report vs. scalp; 4) Surgical Approach: ablation of the cochlea in the present report vs. keeping the cochlea intact; 5) Anesthetics and medications: isoflurane vs. ketamine/xylazine, and the use of pre-anesthetics medications in the present report, such as opioids, i.e., buprenorphine, and mAChR antagonists, such as atropine, which may alter primary afferent or even efferent neuron sensitivity. Among all of these differences, a theoretical model of mechanical activation of the utricle by BCV ^50^ suggests the primary determinant of acceleration vs. jerk sensitivity is the frequency content of the stimulus relative to the major corner frequency of the otolith organ in the direction stimulated. Stimuli below the corner are predicted to show jerk sensitivity, while stimuli near the corner are expected to show acceleration sensitivity. Therefore, differences between species in size of the utricle and differences between stimuli likely explain jerk vs. acceleration scaling of the vCAP. A broad-band stimulus would be expected to evoke more complex vCAPs that do not clearly scale with jerk or acceleration. For this reason, we use the term vCAP for compound action potentials evoked by any vestibular stimulus and reserve VsEP for vCAPs that scale with linear jerk. Macular velocity was not recorded in previous VsEP experiments but based on the present results we would expect the relationship between vCAP and macular velocity to hold even for stimuli where the VsEP scales with linear jerk.

Chirps are used to evoke cochlear responses in animal models and the clinic, such as the chirp-evoked ABR^53^. Special stimuli have been created to overcome travelling wave delays associated with cochlear mechanics^54^. Recent studies have extended these stimuli to the vestibular system to generate VEMPs^55,56^, however, it is unclear how these relatively complex stimuli evoke synchronous neural responses at the end-organ level. Moreover, many of the stimuli which have translated from the cochlea to the vestibular system have been designed to suit unique features of auditory transduction^57^. Hence, it is not apparent if chirps are well suited for otolithic receptor activation. This work reveals that chirps produce robust macular vibration and sensory vCAPs, providing support for their use as part of the neuro-otology test battery. However, it is important to consider the relevant stimulus characteristics for generating transient vestibular responses. Data shows vCAPs respond to the initial onset or offset of the stimulus waveform, with relevant spectral power below 1kHz (Fig.10 & 11). When the transient onset (or offset) is smoothed by increasing the rise-time, the response drops off abruptly (Fig. 10). These results provide a neurophysiological framework for earlier findings, which reported robust VEMPs in humans evoked by band limited chirps (250-1000Hz), chosen because of the ideal sensitivity range of the otoliths^55,58^. Hence, a band limited chirp accompanied with a short rise-time should be considered when designing specific parameters to generate VsEPs, vCAPs, or VEMPs in the laboratory or clinic. Moreover, as the relevant power spectrum approximates the natural frequency of the guinea pig utricle (~500Hz), these vCAPs scale with temporal bone acceleration, rather than jerk, as predicted by the modelling. This relation will of course change with different stimulus parameters and end-organ properties, as mentioned above.

Vibration and sound are excellent stimuli for activating vestibular reflex responses in the clinic, driven by the irregular primary afferent neurons ^14^. However, precisely how BCV and ACS generate vestibular functional responses at the level of the macula are not well understood. Previous results have demonstrated that there are likely differences in macromechanical activation modes of macular receptors for sinusoidal BCV and ACS across frequencies. That is, the relative phase of the VM and macular velocity evoked by sinusoidal stimuli has been shown to be different for BCV, than for ACS, especially at frequencies beyond 300Hz up to kilohertz^59^. At low frequencies (<300Hz), both BCV and ACS VMs and macular vibration responses are approximately ‘in-phase’. Beyond this frequency, BCV microphonics undergo a complex phase shift (lag) relative to macular vibration, up to 2-3 cycles at 1kHz. By contrast, the relative phase of the ACS microphonic and macular velocity remains relatively flat up to high frequencies. Despite differences in timing and activation for sinusoidal BCV and ACS hair cell and macular responses, it is not clear if differences exist in neural responses evoked by transient BCV and ACS. This is relevant for understanding the generation of the BCV and ACS VEMP to punctate stimuli at the bedside for diagnostic purposes. To examine differences in BCV and ACS neural response generation, simultaneous measures of the vCAP and macular velocity were recorded by pulsatile stimulation across animals. Without significant perilymph in the vestibule, results reveal larger vCAP input-output (vCAP IO) amplitudes and slopes for BCV compared to ACS (Fig. 12). However, further investigation identified this difference as a conductive loss from inadequate ACS stimulus coupling and reduced mechanical sensitivity. Here, discrepancies in vCAP IO slope fell away when recording ACS responses with significant perilymph overlying the macula (Fig. 13D & E). Hence, pulsatile sound and vibration vCAP responses share equivalent IO functions and macular vibration thresholds to brief BCV and ACS stimuli, suggesting analogous mechanical activation modes of their sensory receptors across stimuli.

Simultaneous recordings of macular velocity and vCAP responses provide the ability to quantify the level of epithelial vibration at threshold, and suprathreshold levels such as response saturation. Results indicate that for both vibration and sound stimulation the level of macular vibration for vCAP threshold is <0.3μm/s across animals. Moreover, the level of macular vibration for vCAP IO response saturation is <1μm/s. By comparison, for ACS, the level of stapes vibration for vCAP threshold is over an order of magnitude larger than that of the macula, at ~25μm/s.

To determine how macular vibration is related to MET currents entering sensory hair cells, we compared the VM to the macular velocity and macular displacement for sinusoidal tone bursts. The VM is the voltage modulation in the endolymph relative to reference ground measured adjacent to epithelium and reflects changes in the net MET current entering hair cells caused by hair bundle deflection. Results in Fig. 15 show the VM, and therefore the net MET current, is closely aligned with macular displacement over the entire bandwidth tested. Results are consistent with the hypothesis that hair bundles are deflected primarily by otoconial layer displacement, not velocity, and that hair bundle shear is directly related to the macular displacement measured here using LDV ^60^.

While the magnitude of VMs measuring the net MET currents scaled with macular displacement, the magnitude of vCAPs measuring the action potential synchronization scaled with macular velocity (Fig. 14). This difference highlights rate-sensitive signal processing occurring after the MET current ^61^ manifests primarily as a time derivative in sensitive calyx bearing afferents that synchronize to transient stimuli.

## Conclusion

This work sought to examine the relationship between macular macromechanics and action potential generation from irregular striolar afferents to improve our understanding of their stimulation sensitivity and tuning to clinical stimulation modes. Unlike previous studies, which characterized the operation of vestibular primary afferents relative to intense cranial acceleration, this work goes one step further and characterizes synchronous vestibular afferent responses (vCAPs) relative to macular epithelial vibration, VMs and their input drives. Results demonstrate that vCAPs increase in proportion to macular velocity for both ACS and BCV, suggesting the mechanical mechanism of MET activation is the same for both stimuli. In contrast to vCAPs, VMs increased in proportion to macular displacement, indicating that the net MET current entering all hair cells was gated primarily by displacement, not velocity. The difference between VM and vCAP dynamics reflects adaptation signal processing interposed between the MET current and action potential generation in sensitive vestibular afferents ^25,61^, and is the same process responsible for phase-locking of utricular afferent action potentials to audio frequency stimuli ^34^. For brief BCV pulses (<1ms) used in the present study, macular velocities and vCAPs both increased in proportion to temporal bone acceleration. At longer duration BCV pulses, vCAPs increased in proportion to temporal bone jerk, which aligns with previous VsEPs measurements in rodents at lower stimulus frequencies and higher stimulus strengths relative to the present study ^52^.

## Acknowledgments

*This work was supported by a Macquarie University Research Fellowship (MQRF) (CJP), and a NIH DC 006685 (RDR).*

